# Thiomorpholino antisense oligonucleotides inhibit telomerase and limit cancer cell proliferation

**DOI:** 10.64898/2026.09.28.754723

**Authors:** Julieta Rivosecchi, Arthur J. Zaug, Tomáš Lášek, Balázs Schäfer, Marvin H. Caruthers, Thomas R. Cech

## Abstract

Reactivation of telomerase confers immortality to approximately 90% of human tumors by enabling continuous elongation of the DNA at chromosome ends, or telomeres. The telomerase catalytic subunit TERT adds TTAGGG repeats using a portion of the telomerase RNA component hTR as a template. Because telomerase is inactive in most normal somatic cells, it remains an attractive therapeutic target; however, no telomerase inhibitor has yet demonstrated robust clinical efficacy with acceptable safety. Here we evaluate thiomorpholino oligonucleotides (TMOs) as a new class of antisense oligonucleotides targeting the template region of hTR. TMOs incorporate morpholino rings and phosphorothioate linkages, which enhance nuclease resistance, RNA binding and nuclear uptake. Two anti-hTR TMOs inhibited telomerase activity *in vitro* with an IC_50_ below 1 nM, whereas two control TMOs were at least 100-fold less active. HeLa cells treated with anti-hTR TMOs showed progressive telomere shortening, detectable after one week of treatment. Growth inhibition was observed after substantial telomere erosion, and both telomere length and proliferation recovered upon withdrawal of TMOs. These findings establish TMOs as a promising new chemistry for telomerase-targeted therapeutics.

## INTRODUCTION

Telomeric DNA consists of arrays of TTAGGG repeats that progressively shorten with each cell division in somatic cells. Telomere erosion is associated with limited cellular proliferation and replicative senescence in human cells (Harley et al. 1990; Kaul et al. 2011; Schmidt et al. 2024) (See also Passanisi and Spencer 2026). Most cancers circumvent this barrier by reactivating telomerase, which maintains telomeres and supports continued cell division (Kim et al. 1994). Telomerase is a ribonucleoprotein (RNP) reverse transcriptase that counteracts telomere shortening by synthesizing telomeric DNA repeats at chromosome ends using the telomerase RNA component hTR (Greider and Blackburn 1989; Lingner et al. 1997). Because telomerase activity is largely restricted to malignant cells, it has long been considered an attractive target for cancer therapeutics (Gao and Pickett 2022).

The hTR RNA contains an 11-nucleotide sequence complementary to the telomere repeat that base-pairs with the single-stranded telomeric DNA and serves as a template for telomere extension, and additional stem-loop elements that recruit TERT, four H/ACA ribonucleoproteins (the dyskerin complex), and TCAB1 into the RNP complex (Venteicher et al. 2009; Egan and Collins 2010; Rivosecchi et al. 2026). hTR is essential for telomerase activity (Weinrich et al. 1997) and its template is necessarily accessible to nucleic-acid binding, making it a plausible target for antisense oligonucleotide (ASO)-based inhibition (Fig. 1A). Early studies using synthetic oligonucleotides demonstrated that direct hybridization to the hTR template can inhibit telomerase by blocking its binding to telomeres and causing telomeres to shorten (Norton et al. 1996; Shea-Herbert et al. 2002; Chen et al. 2002). Among these oligonucleotides, only Imetelstat, a lipid-conjugated thiophosphoramidate (Asai et al. 2003), has received FDA approval for treatment of a subset of myelodysplastic syndromes (Lennox et al. 2024). However, potential off-target effects may contribute to the hematopoietic toxicity associated with this drug (Kim et al. 2025). Moreover, Imetelstat’s mechanism of action *in vivo* has been revisited; the drug induces ferroptotic cell death, and a mismatched version lacking ASO activity has equivalent anti-proliferative effect in acute myeloid leukemia (AML) cell lines and reduces AML burden in a mouse xenograft model (Bruedigam et al. 2024). Thus, the efficacy of ASOs targeting the template of hTR for inhibiting cancer cell proliferation is an open question.

**FIGURE 1.**
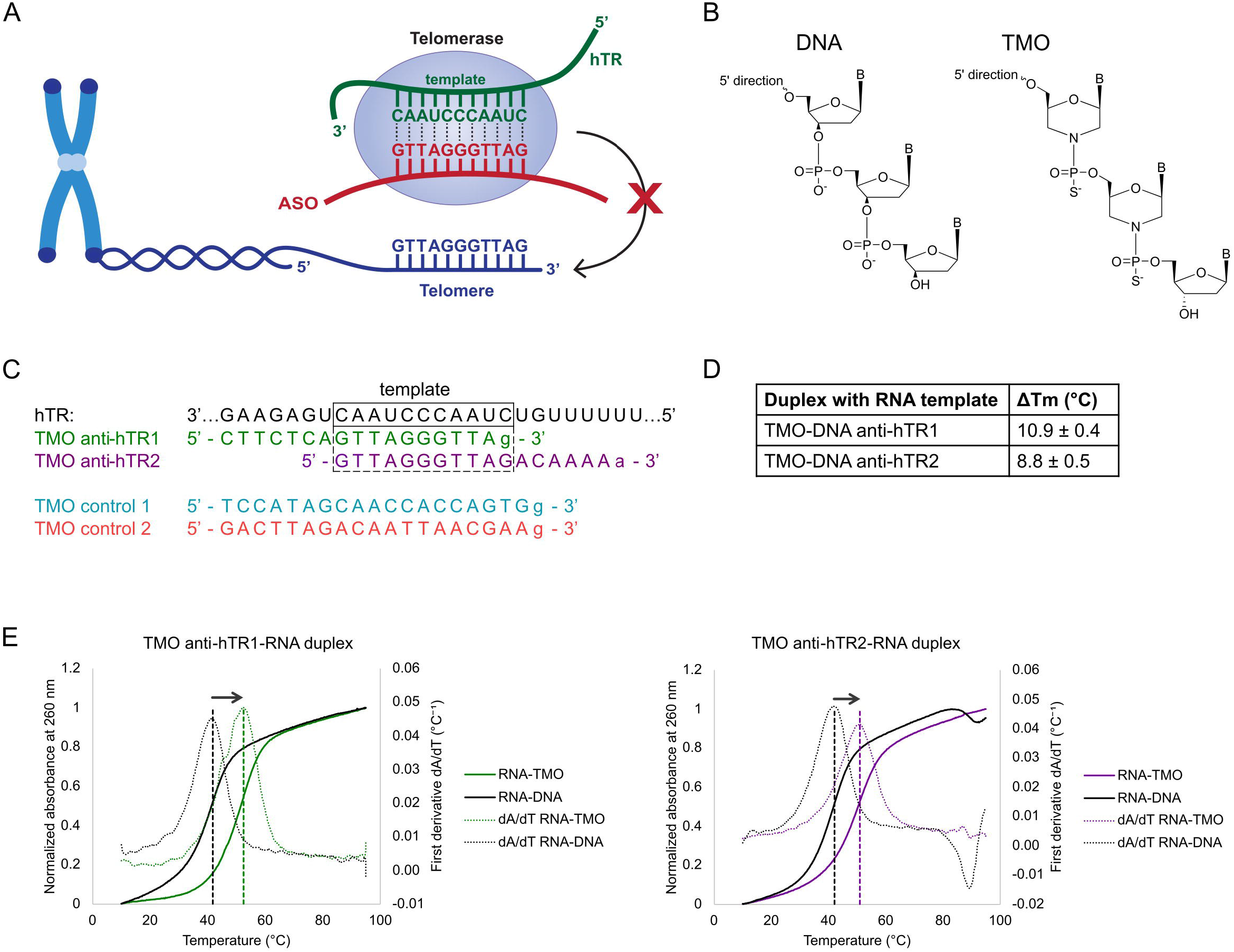
Anti-hTR TMOs bind the hTR template with higher affinity than DNA oligonucleotides. **A.** Antisense oligonucleotides (ASOs) targeting the template region of hTR block its binding to telomeric DNA. Figure created with Adobe Illustrator. **B.** Chemical structures of DNA and thiomorpholino oligonucleotides (TMOs). **C.** Sequences of the hTR template region, anti-hTR TMOs and control TMOs. Capital letters indicate TMO nucleotides, lowercase letters indicate deoxynucleotides, which are a requirement for solid-phase synthesis. **D.** Changes in melting temperature (ΔTm) between TMO-RNA and DNA-RNA duplexes formed with the complementary 11-nt hTR template RNA. Values represent the mean ± standard deviation (SD) of 3 replicates. **E.** Representative UV melting profiles of anti-hTR TMOs and corresponding DNA oligonucleotides duplexed with the complementary 11-nt hTR template RNA, with Tm determined as the peak of the first derivative. Data represent the mean of 3 measurements.

Thiomorpholino oligonucleotides (TMOs) represent a newer class of ASOs that combine phosphorothioate and morpholino chemistries, enhancing stability, RNA binding affinity and nuclear localization (Langner et al. 2020). TMOs have been successfully applied across diverse RNA-targeting strategies, including blocking mRNA splicing (Dumbović et al. 2021), inducing splice-switching (Han et al. 2026), promoting exon-skipping (Dumbović et al. 2021; Le et al. 2022), and functioning as gapmers to elicit RNase-H-mediated degradation of target transcripts (Mejzini et al. 2024).

In this study, we investigated TMOs designed to hybridize to the 11-nucleotide template region of hTR. We show that TMOs bind hTR with higher affinity than DNA oligonucleotides of identical sequence, inhibit telomerase with nanomolar potency *in vitro*, and induce telomere shortening and reduced cell proliferation in telomerase-positive cancer cells. TMO-mediated telomerase inhibition is fully reversible, with telomere elongation and proliferation restored upon withdrawal. These findings position TMOs as a potential strategy for selective telomerase inhibition and establish a foundation for developing next-generation RNA-targeting therapeutics for telomerase-positive cancers.

## RESULTS

### TMOs bind hTR with higher affinity than DNA oligonucleotides

We synthesized two 18-mer TMOs targeting hTR (Fig. 1B) using a modified version of the previously described solid-phase synthesis procedure (Langner et al. 2020). Anti-hTR1 and anti-hTR2 were designed to hybridize to the 11-nt hTR template region, each incorporating seven additional nucleotides extending either 5′ or 3′ from the template, respectively (Fig. 1C). This length was chosen to reduce off-target effects, as the theoretical probability that ASOs longer than 16-mer have a match in the human mRNA sequences is less than 5% (Yoshida et al. 2018). In support of the choice of an 18 nt length, ASO drugs FDA-approved through 2024 that act either by eliciting RNase-H degradation or by blocking RNA splicing range from 18 to 30 nucleotides (Sang et al. 2024). Two control TMOs were synthesized to test for nonspecific cellular effects associated with TMO uptake (Fig. 1C). Purity and molecular weight of all TMOs were verified by liquid chromatography-mass spectrometry (LC-MS) (Supplemental Fig. S1A-E).

To evaluate the thermal stability of anti-hTR TMOs hybridized to RNA, we performed UV thermal denaturation under physiological buffer conditions. Duplexes formed between anti-hTR1 or anti-hTR2 TMOs and an 11-mer RNA corresponding to the hTR template sequence exhibited Tm values that were 10.9°C and 8.8°C higher, respectively, than those of the corresponding DNA-RNA duplexes, indicating that TMOs bind hTR template RNA with higher affinity than DNA oligonucleotides of the same sequence (Fig. 1D-E). Similar results were obtained using a 24-mer RNA containing the hTR template region and flanking sequences, for which TMO-RNA duplexes also displayed higher Tm values than the corresponding DNA-RNA duplexes (Supplemental Fig. S1F-G). These findings are consistent with previous reports (Langner et al. 2020) and demonstrate that TMOs retain enhanced RNA affinity towards the hTR template sequence under physiological buffer conditions.

### TMOs inhibit telomerase activity *in vitro*

We next evaluated the impact of anti-hTR TMOs on telomerase activity using a direct telomerase assay, in which recombinant human telomerase overexpressed in human cells and immunopurified is used to extend a telomeric DNA oligonucleotide with radiolabeled dNTPs. Anti-hTR TMOs inhibited telomerase activity in a dose-dependent manner (Fig. 2A-B), with IC_50_ values of ∼0.5 nM (Fig. 2C), whereas the control TMOs were at least 100-fold less potent. Single-stranded DNA oligonucleotides of identical length and sequence as the TMOs or with the sequence of Imetelstat were >400-fold less inhibitory (Fig. 2A and D), highlighting the efficacy of TMO chemistry. Anti-hTR TMOs showed stronger inhibition than the small-molecule telomerase inhibitor BIBR1532 (Supplemental Fig. S2A). Preincubation of telomerase with anti-hTR TMOs did not enhance telomerase inhibition, suggesting that TMO binding to hTR reaches equilibrium very quickly, < ∼1 minute (Supplemental Fig. S2B).

**FIGURE 2.**
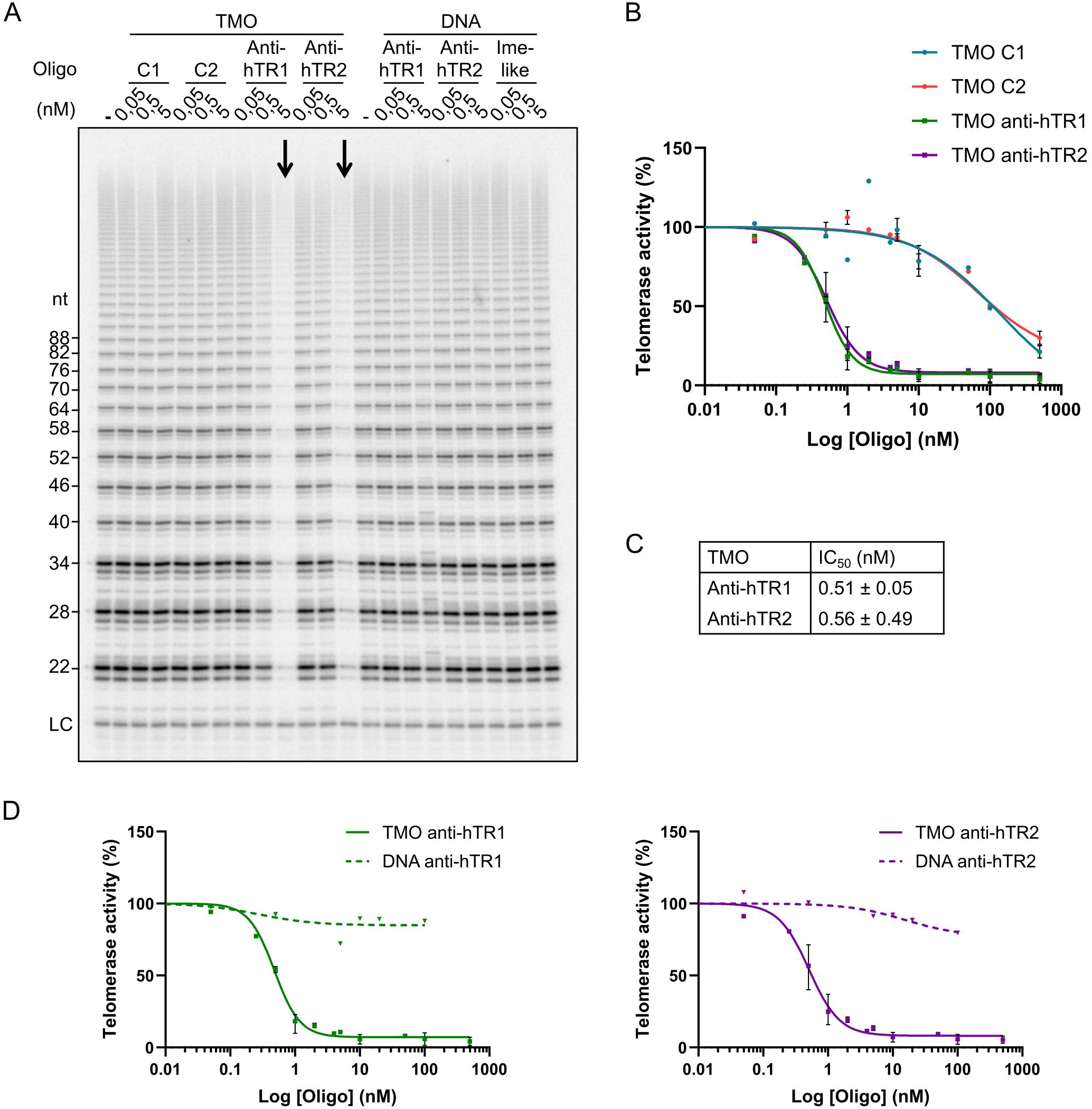
Anti-hTR TMOs inhibit telomerase activity *in vitro*. **A.** Direct telomerase activity assay in the presence of the indicated TMOs and DNA oligonucleotides. Ime-like: DNA oligonucleotide containing the Imetelstat sequence. LC: 18-mer labeled oligonucleotide loading control. The experiment was repeated five times with similar results. **B.** Quantification of telomerase activity for the TMOs shown in panel A. Points shown with error bars represent the mean ± range of 2 or 3 replicates. **C.** Half-maximal inhibitory concentration (IC_50_) values for anti-hTR TMO 1 (mean ± SD of 3 replicates) and TMO 2 (mean ± range of 2 replicates). **D.** Quantification of telomerase activity for anti-hTR TMOs and DNA oligonucleotides including experiment of panel A.

Anti-hTR TMOs might inhibit telomerase either by binding to the hTR template or, given the extended 3’ and 5’ regions of complementarity, by binding beyond the template region and disrupting the RNP complex. To distinguish between these possibilities, telomerase extracts were incubated with TMOs or the corresponding DNA oligonucleotides, separated by native polyacrylamide gel electrophoresis, and analyzed for the presence of free hTR by northern blotting. TMO binding to hTR did not disrupt telomerase RNP integrity, suggesting that anti-hTR TMOs inhibit telomerase by preventing engagement with the telomeric DNA substrate rather than disassembling the enzyme complex (Supplemental Fig. S3).

### TMOs induce telomere shortening in HeLa cancer cells

To investigate the cellular consequences of anti-hTR TMO treatment, we used a HeLa clone with relatively short telomeres previously generated in our laboratory (Wu et al. 2025) to reduce the time required for telomeres to reach a critically short length. We first assessed nuclear uptake in this HeLa cell line using a FAM-conjugated TMO (TMO-FAM). After 48 h of incubation with 50-200 nM TMO-FAM, we observed nuclear fluorescence in less than 2.5% of cells through spontaneous uptake. In contrast, Lipofectamine 3000-mediated transfection resulted in efficient nuclear localization of TMO-FAM (Supplemental Fig. S4A). All subsequent cellular experiments were performed using Lipofectamine-mediated TMO transfection. To minimize toxicity while maintaining efficient nuclear delivery, we selected a TMO concentration of 50 nM (100-fold above the IC_50_ determined *in vitro*), which resulted in nuclear localization in 72% of cells with minimal cytotoxicity after 48 h of incubation. Given that TMOs are stable inside cells for at least 5 days (Le et al. 2022), cells were transfected twice weekly (Supplemental Fig. S4B).

Cells were treated with control TMOs, anti-hTR TMOs or 10 µM of the telomerase inhibitor BIBR1532 as a positive control. Telomere restriction fragment (TRF) analysis was performed periodically to monitor changes in telomere length. Our results showed that untreated HeLa cells displayed an average telomere length of approximately 4 kb, whereas BIBR1532 treatment induced shortening to ∼3 kb after 8 weeks, consistent with prior studies (Nakashima et al. 2013). Anti-hTR TMOs caused progressive telomere erosion, reaching ∼3 kb and ∼2.7 kb after 8 weeks of treatment with anti-hTR1 and anti-hTR2, respectively. In contrast, cells treated with control TMOs unexpectedly exhibited modest telomere lengthening (Fig. 3A-B).

**FIGURE 3.**
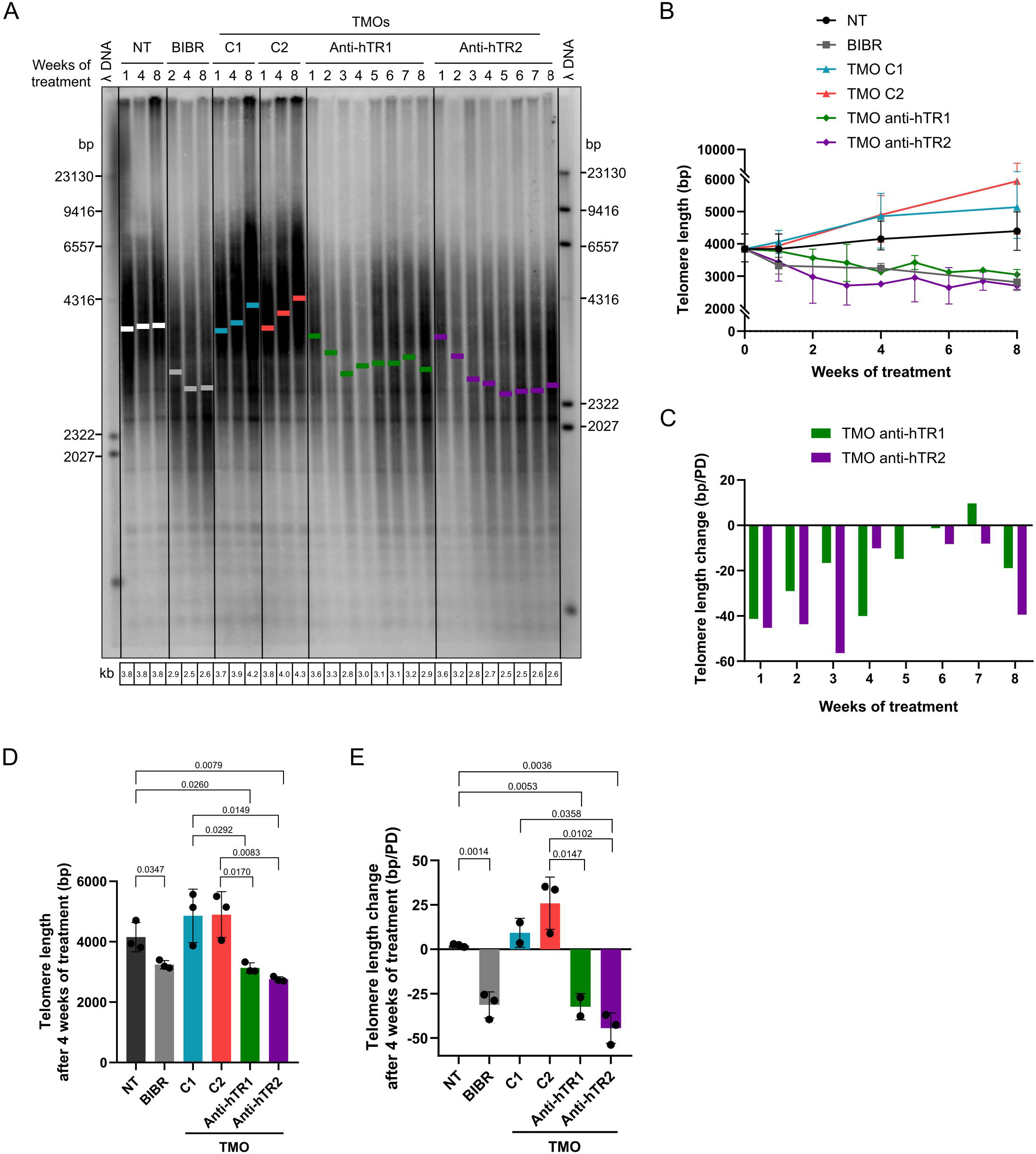
Anti-hTR TMOs induce telomere shortening in telomerase-positive cancer cells. **A.** Terminal restriction fragments (TRF) analysis of genomic DNA extracted from untreated (NT) HeLa cells, or cells treated with BIBR1532 (BIBR) or the indicated TMOs. The experiment was repeated twice for anti-hTR1 and three times for anti-hTR2 TMOs, with similar results. **B.** Quantification of TRF values shown in panel A. Each data point represents the mean of 2 or 3 biological replicates ± range. **C.** Rate of telomere loss or gain per week, expressed in base pairs (bp) per population doubling (PD), average of 2 biological replicates. **D.** Quantification of telomere length from TRF analysis 4 weeks after treatment with BIBR, TMOs or no treatment, n = 3 biological replicates. *p*-values were determined using an unpaired t-test. **E.** Telomere length changes 4 weeks after treatment with BIBR, TMOs or no treatment, expressed in bp/PD, n = 3 biological replicates. *p*-values were determined using an unpaired t-test.

Telomere shortening occurred most rapidly during the first 3 weeks of anti-hTR TMO treatment and reached a plateau by week 4 (Fig. 3C). By this time, telomeres were significantly shorter in anti-hTR TMO-treated cells compared with untreated or control-treated cells (Fig. 3D). Moreover, anti-hTR1 induced ∼790 bp loss over 4 weeks, and anti-hTR2 induced ∼1200 bp loss, corresponding to ∼32 bp and ∼44 bp per population doubling (PD), respectively (Fig. 3E). These rates slightly exceeded the ∼31 bp/PD observed with BIBR1532 and previously reported for telomerase-positive cancer cell lines (Nakashima et al. 2013; Damm et al. 2001). Together, these results indicate that TMOs are effectively internalized into the nucleus of HeLa cells and induce rapid telomere shortening of telomeres beginning within 1 week of treatment.

### Effect of TMOs on cancer cell proliferation

To assess the effect of anti-hTR TMOs on cancer cell growth, HeLa cells were replated weekly to maintain log-phase conditions, and cumulative PD and growth rate were calculated. BIBR1532 treatment modestly reduced proliferation over 10 weeks, compared with untreated cells (Fig. 4A and Supplemental Fig. S4C), consistent with previous reports (Nakashima et al. 2013). Transfection of control TMOs caused a mild, stable decrease in proliferation relative to untreated cells (Fig. 4A and Supplemental Fig. S4C). Both anti-hTR TMOs progressively reduced proliferation across the 10-week experiment compared with untreated and control-TMO treated cells, as shown by PDs (Fig. 4A) and growth rate (Supplemental Fig. S4C), with variability among biological replicates (Fig. 4A). Notably, growth rate was significantly reduced by week 4, coinciding with shortening of telomeres (Fig. 4B and Supplemental Fig. S4D).

**FIGURE 4.**
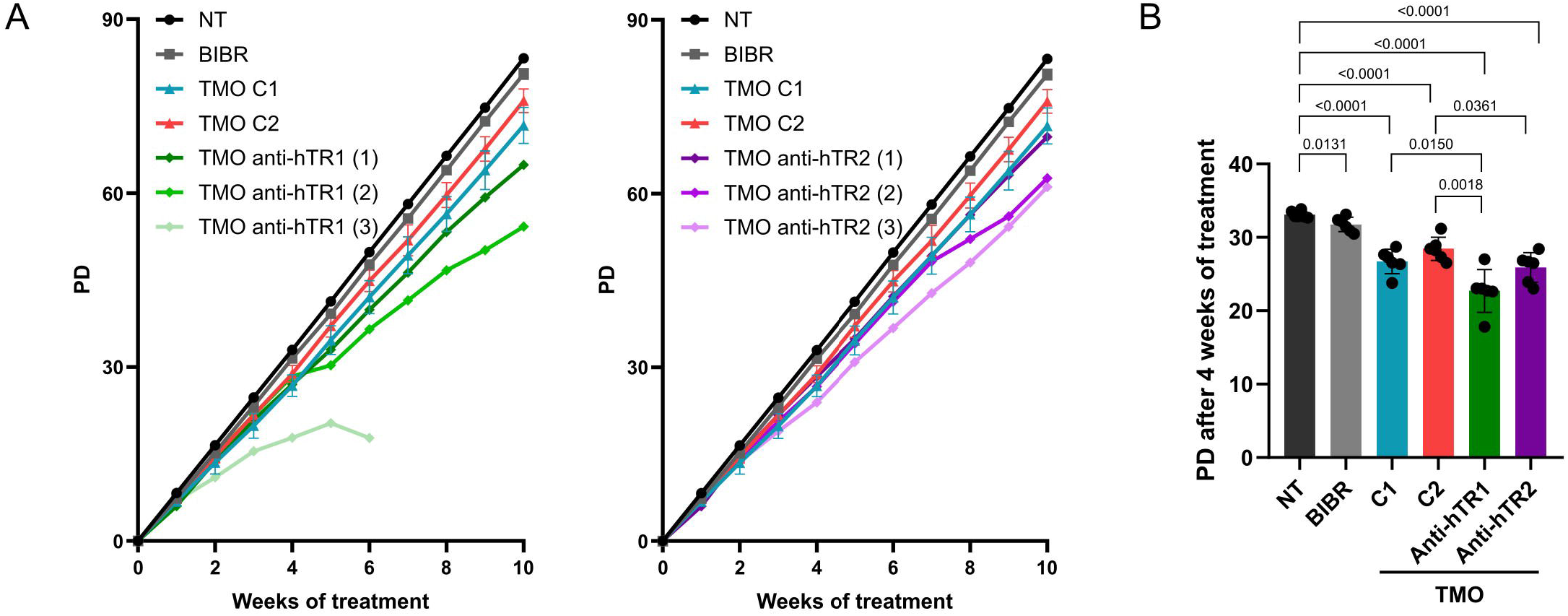
TMOs limit cancer cell proliferation. **A.** Cumulative growth expressed as population doublings (PD) of untreated HeLa cells, treated with BIBR1532 or the indicated TMOs, monitored weekly over 10 weeks. Data for NT and BIBR-treated cells represent the mean ± SD of 3 independent biological replicates and are shown in both panels for comparison; for anti-hTR1 (left panel) and anti-hTR2 TMO-treated cells (right panel), the three independent biological replicates (1-2-3) are shown individually. **B.** Cumulative growth (PD) of untreated HeLa cells, 4 weeks after treatment with BIBR1532 or the indicated TMOs. Data represent the mean ± SD of six independent biological replicates. *p*-values were determined using an unpaired t-test.

As a specificity control, we treated telomerase-negative U2OS cells, which rely on the Alternative Lengthening of Telomeres (ALT) pathway for the maintenance of telomeres (Bryan et al. 1997). Lipofectamine LTX transfection of TMOs at 50 nM resulted in 55% TMO-FAM-positive cells after 24 h of incubation (Supplemental Fig. S5A). TMOs were transfected twice weekly, as in HeLa cells. Our results showed that U2OS cell proliferation was not affected by anti-hTR TMOs at 50 nM (Supplemental Fig. S5B) or 75 nM (Supplemental Fig. S5C). Moreover, anti-hTR TMOs did not alter telomere length after 4 weeks of treatment (Supplemental Fig.S5D), although subtle changes would be difficult to detect given the heterogeneous, long telomeres of U2OS cells. These data are consistent with a functional link between telomerase inhibition, telomere shortening and reduced cell growth induced by TMOs.

### Reversibility of telomerase inhibition by TMOs

Considering our proposed mechanism-of-action of the anti-hTR TMOs, we hypothesized that their effects should be reversible upon cessation of treatment. To assess reversibility, cells treated for 6 weeks were transferred to medium lacking TMOs and monitored for an additional 4 weeks. Remarkably, telomere elongation was evident within 1 week of withdrawing anti-hTR2 TMO (week 7 of the experiment), as mean telomere length increased from 2.1 kb to 2.6 kb, followed by continued extension to 4.1 kb by 4 weeks after removal of treatment (week 10 of the experiment) (Fig. 5A-B). Coincident with telomere length recovery, cell proliferation progressively improved following TMO withdrawal, and after 3 weeks (week 9 of the experiment), growth rate was indistinguishable from untreated control cells (Fig. 5C). These results demonstrate that TMO-mediated telomerase inhibition is fully reversible, as expected for an anti-sense mechanism.

**FIGURE 5.**
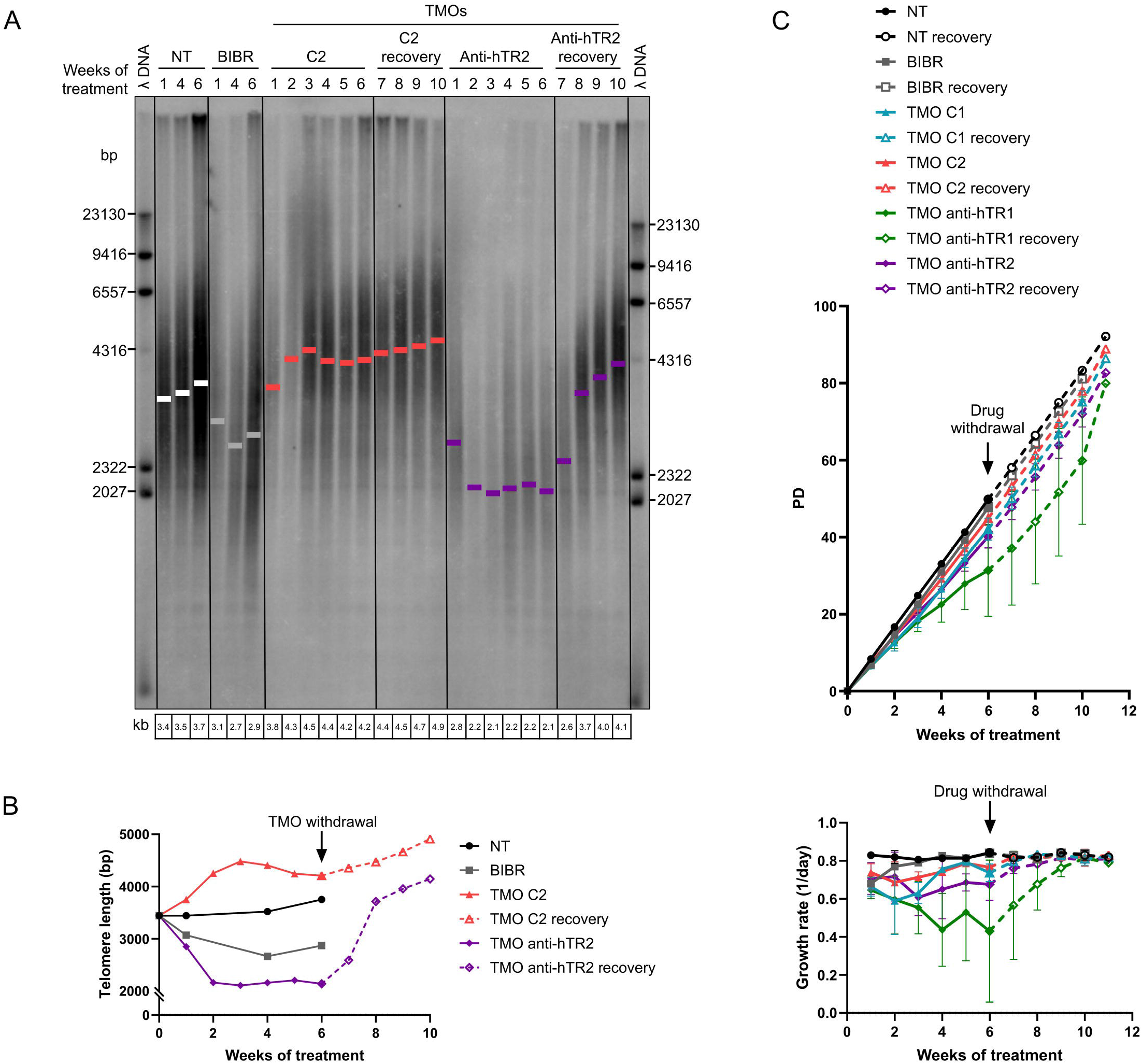
TMO-mediated telomerase inhibition is reversible. **A.** TRF analysis of genomic DNA extracted from untreated HeLa cells, or cells treated with the indicated TMOs. After six weeks of treatment, cells were cultured in normal medium without TMOs for an additional 4 weeks (weeks 7-10). **B.** Quantification of TRF values from panel A. **C.** Growth rate of HeLa cells untreated or treated with BIBR or the indicated TMOs, monitored weekly over six weeks of treatment and five additional weeks after drug withdrawal. Data represent the mean ± SD of 3 independent biological replicates.

## DISCUSSION

Targeting hTR has long been considered an attractive therapeutic strategy for telomerase-positive cancers. However, achieving high affinity, specificity, and cellular efficacy has remained challenging. Here, we show that TMOs directed against the hTR template region of telomerase constitute potent telomerase inhibitors *in vitro* and *in vivo*.

### High RNA affinity enables efficient template-blocking inhibition

Thermal denaturation analyses demonstrated that TMOs form more stable duplexes with hTR than DNA oligonucleotides of identical sequence, increasing the melting temperature by 9-11°C. Although this may seem like a modest increase, note that it is enough to raise the Tm to 50°C, safely above the mammalian temperature of 37°C. This enhanced affinity is particularly relevant for template-blocking strategies, which rely on sustained occupancy of the hTR template to prevent telomeric DNA binding. The increased RNA affinity likely contributes to the sub-nanomolar IC_50_ values observed in direct telomerase assays. We found that the TMOs did not disrupt telomerase RNP integrity. Instead, TMOs inhibit telomerase by directly occluding the RNA template while preserving the overall RNP structure.

### Unprecedented telomerase-inhibitory potential of TMOs

Anti-hTR TMOs inhibited telomerase activity with an IC_50_ of approximately 0.5 nM, regardless of the presence of extra nucleotides at either the 5′ or 3′ end of the oligonucleotide. This potency is unprecedented, as previously reported inhibitors, including Imetelstat and BIBR1532, have IC_50_ values at least 20-fold higher (Damm et al. 2001; Marian et al. 2010). In cancer cells, this strong *in vitro* efficacy translated into an immediate inhibition of telomerase and rapid telomere shortening, detectable after only one week of treatment. Although this rapid and potent cellular response is therapeutically attractive, its reversibility indicates that continuous TMO exposure is required to maintain telomerase inhibition, suggesting that sustained treatment would be necessary for clinical applications. At the same time, the rapid reversibility of the telomere shortening and cell proliferation phenotypes upon TMO withdrawal is an attractive feature for a therapeutic, because if a patient were to suffer from stem-cell depletion upon telomerase inhibition, the drug could be discontinued.

### Rapid telomere shortening and the ∼3 kb threshold

One of the major challenges of telomerase inhibition is the long lag period required until telomeres are sufficiently shortened to impair cell proliferation. Thus, we decided to use a HeLa clone with relatively short telomeres (∼4 kb). Anti-hTR TMOs induced rapid telomere erosion, reaching ∼3 kb by week 4 before stabilizing. Why telomeres failed to shorten further remains unclear. One plausible explanation is that cells harboring telomeres shorter than approximately 3 kb undergo cell death, leading to selective loss of cells with critically short telomeres and resulting in enrichment of cells with lengths near the survival boundary.

Previous studies have shown that telomerase inactivation can promote the emergence of ALT-positive revertants, albeit at low frequency (Bechter et al. 2004; Queisser et al. 2013; Min et al. 2017). Interestingly, in the 10-weeks period of anti-hTR TMO treatment, we failed to detect telomere elongation or discrete bands by TRF, characteristic of ALT telomeres. Likewise, we observed no emergence of rapidly dividing cells following the initial growth inhibition. Thus, our observations are consistent with the anti-hTR TMOs not promoting activation of the ALT pathway under our experimental conditions, although additional ALT hallmarks would need to be assessed to confirm this conclusion.

Unexpectedly, control TMOs or perhaps the lipofection protocol itself induced modest telomere elongation. The mechanism underlying this sequence-independent effect remains unknown and will require further investigation to determine whether control TMOs indirectly influence telomerase recruitment or activity.

### Growth inhibition is telomerase-dependent and reversible

The oligonucleotide Imetelstat, which has a sequence complementary to the hTR template, has recently been reported to induce ferroptotic cell death rather than telomerase-associated cell death (Bruedigam et al. 2024). In contrast, four lines of evidence support TMOs acting as anti-sense oligonucleotides in cells: (1) The two control TMOs, which had the same chemistry as the anti-hTR TMOs, did not elicit telomere shortening. (2) Telomere shortening was detectable only after one week of TMO treatment and clearly preceded the impairment of cell proliferation, supporting a causal relationship between telomerase inhibition, telomere erosion, and growth suppression. (3) Following TMO withdrawal, telomeres rapidly elongated and cell proliferation recovered, further demonstrating that these effects are linked. (4) Telomerase-negative U2OS cells showed no changes in telomere length or proliferation when treated with anti-hTR TMOs, reinforcing the specificity of the response. Although control TMOs modestly reduced proliferation in both HeLa and U2OS cells, these effects were uncoupled from telomere dynamics and likely reflect sequence-independent cellular stress.

### Limitations of the study

This study has several limitations that should be addressed in the future. First, all cellular experiments were performed in HeLa cells. Although HeLa cells are a standard model for telomere and telomerase biology, response to TMOs might be stronger or weaker in other telomerase-positive cancer types. Validation in additional cancer models will be essential to establish the generality of these findings.

Second, TMOs were delivered exclusively by lipofection. While suitable for mechanistic studies, this transfection is not clinically relevant and can introduce sequence-independent toxicity (Lv et al. 2006), as reflected in the mild proliferation defects observed with control TMOs. Future studies will require evaluation of lipid nanoparticle delivery and potentially free delivery of conjugated TMOs.

Third, cellular experiments were conducted at a single TMO concentration selected to balance nuclear uptake and toxicity. Dose-response relationships for telomere shortening, proliferation, and reversibility remain to be defined. Determining whether lower doses can sustain telomerase inhibition, or whether higher doses introduce off-target effects, will be important for therapeutic development.

Fourth, telomere length measurements relied on TRF analysis, which provides population-average telomere lengths but cannot detect critically short telomeres within individual cells. Long-read sequencing (Sanchez et al. 2024; Karimian et al. 2024; Schmidt et al. 2024) would help clarify whether TMOs preferentially affect specific telomere subsets or accelerate the emergence of critically short telomeres.

### In conclusion

Together, these results indicate that TMOs are a promising chemistry for telomerase inhibition. Given that Imetelstat is no longer thought to act *in vivo* through an anti-telomerase mechanism, these TMOs appear to represent the first successful ASOs targeting telomerase. Their high affinity for hTR, potent and reversible suppression of telomerase activity, and rapid induction of telomere shortening position TMOs as a promising class of antisense therapeutics for telomerase-positive cancers. Future work should focus on optimizing delivery strategies beyond lipofection and evaluating TMO efficacy across diverse cancer models.

## MATERIALS AND METHODS

### Oligonucleotide synthesis

TMOs and control DNA oligonucleotides were synthesized on solid support on an Applied Biosystems 394 DNA/RNA synthesizer. The TMO sequences were prepared according to a modified method (unpublished) of the previously reported procedure (Langner et al. 2020). DNA phosphoramidites were coupled under standard conditions. Following each coupling step, the nascent P(III) linkage was converted to the corresponding P(V) thiophosphoramidate (TMO) or phosphorothioate (DNA) linkage via sulfurization (DDTT, 0.05 M). Unreacted hydroxyl groups were capped using standard Cap A (acetic anhydride) and Cap B (1-methylimidazole) solutions. Detritylation was performed using 3% trichloroacetic acid in dichloromethane. After completion of synthesis, oligonucleotides were cleaved from the solid support and deprotected using aqueous ammonia (28%) at 55 °C for 16 h. Crude oligonucleotides were purified by ion-pair reversed-phase HPLC (IP-RP-HPLC), in an Agilent 1200 series HPLC system with a XBridge Oligonucleotide BEH C18 column. The final products were converted into sodium salts by dissolving them in 0.3 M sodium acetate buffer (pH 7.5), followed by the removal of excess salts from the solutions via centrifugation using an Amicon Ultra centrifugal filter (3 kDa molecular weight cutoff).

Quality of the oligonucleotides was assessed by liquid chromatography-mass spectrometry (LC–MS). An Agilent Technologies LCMS-QTOF (Agilent 6530 series Q-TOF LC/MS spectrometer) was used for the purification by the application of an ACQUITY UPLC BEH C18 Column (130Å, 1.7 μm, 2.1 mm X 100 mm). The buffers were A (950 ml of water, 25 ml of methanol (MeOH), 2.5 ml of triethylamine (TEA), 26 ml of hexafluoro-2-propanol (HFIP)); B (50 ml of water, 925 ml of MeOH, 2.5 ml of TEA, 26 ml of HFIP) with the flow of 0.2 ml/min; Gradient: A (0-5 min), A→B (5-30 min), B (30-35 min), B→A (35-36 min), A (36-45 min). The LC/MS spectra were recorded in negative mode, the molecular weights were calculated by subtraction of the charge from the formula weight (F_W_), divided by the charge, calculated and observed mass-to-charge (m/z) ratios. (Formula weight (F_W_) = observed weight −3 charge/3 charge).

Two 20-mer control TMOs were synthesized to test for effects of the TMO chemistry and the lipofection on cell growth: TMO control 1 (C1), which targets exon 4 of the non-essential Integrin Subunit Alpha 4 (ITGA4) gene, and a nontargeting TMO control 2 (C2) validated in previous studies (Dumbović et al. 2021).

### UV thermal denaturation

Thermal denaturation experiments were performed on a Cary 100 Bio UV–Vis spectrophotometer equipped with a temperature controller, with absorbance monitored at 260 nm. Duplexes were prepared by mixing RNA with the corresponding TMO or DNA oligonucleotide in a 1:1 molar ratio (1.0 µM of each strand) in Mg^2+-^containing buffer (10 mM potassium phosphate, 100 mM KCl, 1 mM MgCl_2_, pH 7.0). For each duplex, a 3 mL master mix was prepared and split into three quartz cuvettes (1 cm path length), which were measured in parallel as triplicates together with the corresponding buffer blank. Samples were heated from 25 to 95 °C at 10 °C min^−1^ and held at 95 °C for 5 min. They were then cooled to 10 °C at 0.5 °C min^−1^, held at 10 °C for 5 min, and heated from 10 to 95 °C at 0.5 °C min^−1^. Absorbance was recorded at 0.2 °C intervals during both slow temperature ramps. All reported Tm values were derived from the final heating ramp (10 → 95 °C). For Tm determination, each melting trace was corrected individually by subtracting the buffer blank measured under identical conditions. The temperatures recorded in the individual cell positions differed slightly, so at each acquisition point the temperature used for the final analysis was the mean of the 3 replicate temperatures. The blank-corrected absorbances of the 3 replicates were averaged to give the final melting curve (426 data points between 10 and 95 °C). The curve was then normalized as Anorm = (A − Amin)/(Amax − Amin). Tm was defined as the temperature at the maximum of the first derivative, dAnorm/dT. Numerical differentiation strongly amplifies experimental noise, so the derivative was obtained by local polynomial (Savitzky–Golay-type) regression rather than by simple finite differences. At each data point, a third-order polynomial was least-squares fitted to the 31 surrounding points (the point itself ± 15 neighbors, a window of ≈6 °C). The slope of the fitted polynomial at that point was taken as dAnorm/dT. At the ends of the temperature range, the window was truncated to the available points. The fits were performed directly on the measured temperature axis, so no interpolation or resampling was required. The Tm values were robust to the choice of window: windows between 21 and 41 points changed individual Tm values by less than 0.5 °C. They also agreed within 0.5 °C with Tm values obtained by resampling to a 1 °C grid followed by a 5-point Savitzky–Golay derivative. We therefore estimate the uncertainty of Tm as ±0.5 °C. The effect of the modification on duplex stability was expressed as ΔTm = Tm (RNA/TMO) – Tm (RNA/DNA). Total hyperchromicity was calculated as the ratio of the mean absorbance at the high-temperature end (last ten data points, ≈93–95 °C) to that at the low-temperature end (first five data points, ≈10–11 °C).

### Purification of human telomerase

HEK293T/17 (ATCC CRL-11268) cells were transfected with plasmids containing hTERT (pvan107-3xFLAG) and hTR (pSUPER-hTR) at a 1:3 molar ratio using Lipofectamine 2000. The cells were further expanded 3-fold 24 h after transfection and then harvested 24 h later. To purify telomerase, the cell pellet was lysed with CHAPS lysis buffer (10 mM Tris–HCl pH 7.5, 1 mM MgCl_2_, 1 mM EGTA, 0.5% CHAPS, 10% glycerol, 5 mM beta-mercaptoethanol) for 45 min at 4 °C on a rotator. The lysate was then clarified by centrifugation at 13,000 x g at 4 °C for 15 min. Anti-FLAG resin (A2220, Sigma-Aldrich) was added to the clarified supernatant and the samples incubated on a rotator or for 4 h (overnight) at 4°C. The anti-FLAG resin was washed 3x with wash buffer (20 mM HEPES–NaOH pH 8.0, 2 mM MgCl_2_, 0.2 mM EGTA, 0.1% NP-40, 10% glycerol, 1 mM DTT) before elution using wash buffer supplemented with 0.25 mg/ml 3xFLAG peptide (F4799, Sigma-Aldrich). Purified telomerase complex was verified by western blotting with anti-FLAG antibody.

### Telomerase activity assay

Activity of the immunopurified human telomerase complex was determined by a direct assay modified from a published protocol (Forino et al. 2025). The reaction mixture (20 µL) contained 1x human telomerase assay buffer (50 mM Tris-HCl at pH 8.0, 50 mM KCl, 50 mM NaCl, 1 mM MgCl_2_, 5 mM 2-mercaptoethanol, 1 mM spermidine), 0.05 µM telomeric DNA primer, 0.5 mM dTTP, 3 µM dGTP, 10 µM dATP, 0.17 µM ^32^P-dATP (3000 Ci/mmol, Perkin Elmer) and the corresponding compounds (BIBR, TMOs or DNA oligonucleotides). Following a 1 h incubation at 30 °C, reactions were stopped with the addition of 100 µL of 3.6 M NH_4_OAc containing 20 µg of glycogen. Ethanol (500 µL) was added for precipitation. After incubating overnight at −80 °C, samples were centrifuged for 15 min at 4 °C. Pellets were washed with 70% ethanol and resuspended in 10 µL of H_2_O followed by 10 μL of 2x loading buffer (94% formamide, 0.1x TBE, 0.1% bromophenol blue, 0.1% xylene cyanol). The heat-denatured samples were loaded onto a 10% polyacrylamide/7 M urea/1x TBE gel for electrophoresis. After electrophoresis, the gel was dried and quantified by using an Amersham Typhoon (Cytiva).

### Incubation of purified telomerase with oligonucleotides for analysis of RNP integrity

Purified telomerase was incubated with TMO or DNA oligonucleotides (5 nM) in 1x human telomerase assay buffer (50 mM Tris-HCl at pH 8.0, 50 mM KCl, 50 mM NaCl, 1 mM MgCl_2_, 5 mM 2-mercaptoethanol, 1 mM spermidine) for 15 min at 30 °C. As a positive control for the release of hTR from the RNP complex, purified telomerase was incubated with 5 µl of proteinase K (20 mg/ml) in proteinase K buffer (20 mM Tris pH7.5, 5 mM CaCl_2_) for 1 h at 37°C before analysis on the native gel.

### hTR northern blot

Telomerase incubated with oligonucleotides or not treated was separated on a native 5% polyacrylamide/1xTBE gel. The gel was run at 35 W at 4°C until the xylene cyanol dye had migrated 10 cm. Nucleic acid was transferred to Hybond-N+ membrane (Cytiva #RPN203B) at 2 amps in 0.5x TBE at 4 °C for 1 h. Nucleic acid was crosslinked to the membrane using a CL-1000 ultraviolet crosslinker (Krackeler Scientific) at 254 nm with 1200×100 µJ/cm^2^. The membrane was blocked using PerfectHyb PLUS (Sigma-Aldrich #H7033) for 30 min at 60 °C. hTR probe (50 pmol) was labeled with 2 μl of ^32^P-γ-ATP (6000 ci/mmol), 1 μl of T4 Polynucleotide Kinase (NEB) and 1x kinase buffer in a 20 μl reaction at 37°C for 1 h, and unincorporated nucleotides were removed with Micro Bio-Spin P-6 Gel Columns (BioRad #7326221). 10^7^ cpm of radiolabeled hTR probe was added and incubated overnight at 60 °C. Membrane was rinsed with 25 ml 2x SSC/0.1% SDS and washed 3x with 75 ml 2x SSC/0.1% SDS for 15 min at 60 °C, then rinsed with 25 ml 0.2x SSC/0.1% SDS and washed with 75 ml 0.2x SSC/0.1% SDS for 15 min. Membrane was wrapped in saran wrap and exposed to a phosphor screen for 1-2 h and analyzed using an Amersham Typhoon (Cytiva).

### Culture, transfection and cell proliferation assays of human cell lines

HeLa FLAG-HaloTag-RTEL1 HA-mEos3.2-FKBP12^F36V^-TRF2 clone 15 (Wu et al. 2025), U2OS (ATCC #HTB-96) and HEK293T (ATCC #CRL-11268) cells were incubated at 37 °C, 5% CO_2_, and ≥80% humidity in DMEM with 10% v/v fetal bovine serum (FBS), 1x GlutaMAX-I and 1% w/v penicillin/streptomycin.

Every Monday, cells were counted and 1 x10^5^ HeLa cells or 1.5 x10^5^ U2OS cells were seeded in 6-well plates. 24 h later, 50 nM of TMOs were transfected for 4 h using Lipofectamine 3000 (Invitrogen #L3000015) and Lipofectamine LTX (Invitrogen #15338100) for HeLa and U2OS, respectively, according to the manufacturer’s protocol. 48 h after transfection, cells were transferred to 10 cm dishes. On Fridays, TMOs were transfected again under the same conditions. BIBR was used at a concentration of 10 μM, and the drug was added every 2-3 days. Population doublings (PDs) were calculated as [log N(t)-log N(t0)]/log 2, where N(t0) and N(t) represent the numbers of cells seeded and counted 7 days later, respectively. Growth rates were calculated as [ln (N(t)/N(t0)]/Δt, where Δt represents the number of days elapsed.

### Telomere restriction fragment (TRF) analysis

Genomic DNA was extracted from cells using the GenElute Mammalian Genomic DNA Kit (Sigma-Aldrich G1N70-1KT) and measured using the Qubit 1X dsDNA HS Assay Kit. DNA (10 μg) was digested at 37 °C overnight with HinfI and RsaI in CutSmart buffer. The digested DNA was separated on a 0.8% agarose gel at 50 V for 18 h. The gel was incubated with 0.25 M HCl for 15 min, 0.5 M NaOH and 1.5 M NaCl for 30 min, then neutralized with Tris-NaCl buffer (1 M Tris, pH 7.5, 1.5 M NaCl) for 30 min. Nucleic acid was transferred to Hybond-N+ membrane (Cytiva #RPN203B) by capillarity overnight with 10X SSC buffer. Nucleic acid was crosslinked to the membrane using a CL-1000 ultraviolet crosslinker (Krackeler Scientific) at 254 nm with 1200×100 µJ/cm^2^. The membrane was blocked using PerfectHyb PLUS (Sigma-Aldrich #H7033) for 30 min at 50 °C. TelC probe (2.5 μM) was labeled with 5 μl of ^32^P-γ-ATP (6000 Ci/mmol), 1 μl of T4 Polynucleotide Kinase (NEB) and 1x kinase buffer in a 20 μl reaction at 37°C for 1 h, and unincorporated nucleotides were removed with Micro Bio-Spin P-6 Gel Columns (cat# 7326221, Bio-Rad). Radiolabeled probe was added and incubated overnight at 50 °C. Membrane was rinsed 3x with 75 ml 0.1x SSC/0.1% SDS and washed 3x with 75 ml 0.1x SSC/0.1% SDS for 15 min at 50 °C. Membrane was wrapped in saran wrap and exposed to a phosphor screen for 72 h and analyzed using an Amersham Typhoon (Cytiva).

## ACKNOWLEDGMENTS

J.R. is supported by a fellowship from the AIRC Foundation for Cancer Research in Italy (Dottor Mauro Fiori ID 31173). T.R.C. is an investigator of the Howard Hughes Medical Institute. T.L., B.S. and M.H.C. acknowledge support from the University of Colorado. We thank Theresa Nahreini, Emily Proksch and the Biochemistry Cell Culture Facility (RRID:SCR_018988) for technical assistance and equipment use during these studies.

---

**SUPPLEMENTAL FIGURE 1. A-D.** Liquid chromatography-mass spectrometry (LC-MS) characterization of TMOs showing UV diode array detector (DAD) chromatograms (upper panels), total ion chromatograms (TICs) (middle panels), and electrospray ionization (ESI) mass spectra (lower panels), confirming the purity and expected molecular weights (MW) of anti-hTR TMOs (**A**-**B**) and control TMOs (**C**-**D**). **E.** Calculated and measured MW of TMOs from ESI panels in A-D. **F.** ΔTm between TMO-RNA and DNA-RNA duplexes formed with the 24-mer hTR RNA containing the template region and flanking sequences. Values represent the mean ± SD of 3 replicates. **G.** Representative UV melting profiles of anti-hTR TMOs and corresponding DNA oligonucleotides duplexed with the complementary 24-mer hTR RNA containing the template region and flanking sequences. Data represent the mean of 3 measurements.

**SUPPLEMENTAL FIGURE 2. A.** Direct telomerase activity assay in the presence of the indicated TMOs, DNA oligonucleotides and BIBR1532. **B.** Direct telomerase activity assay and corresponding quantification in the presence of the indicated TMOs under the indicated preincubation conditions. LC: 18-mer labeled oligonucleotide loading control.

**SUPPLEMENTAL FIGURE 3. A.** Native gel and northern blot analysis of hTR with purified telomerase not treated (NT) or after incubation with the indicated TMO and DNA oligonucleotides, with *in vitro*-transcribed (IVT) hTR showing where released telomerase RNA would run. The experiment was repeated four times with similar results.

**SUPPLEMENTAL FIGURE 4. A.** Quantification of nuclear fluorescence in HeLa cells transfected or not with TMO-FAM at the indicated concentrations. Error bars represent the mean of 2 biological replicates ± range. **B.** Experimental protocol for sustained transfection of HeLa cells with control or anti-hTR TMOs. **C.** Growth rate of untreated HeLa cells, or cells treated with BIBR1532 (BIBR) or TMOs, expressed as day^-1^ (number of generations per day) and calculated weekly. Data represent the mean ± SD of at least three independent biological replicates. **D.** Growth rate of HeLa cells 4 weeks after treatment with BIBR or TMOs, expressed as day^-1^. Data represent the mean ± SD of six independent biological replicates.

**SUPPLEMENTAL FIGURE 5. A.** Quantification of nuclear fluorescence in U2OS cells transfected with TMO-FAM at the indicated concentrations using Lipofectamine LTX. **B.** Cumulative growth (PD) and growth rate (PD day^-1^) of U2OS cells untreated or treated with BIBR or 50 nM of TMOs, monitored weekly over 4 weeks. **C.** Cumulative growth (PD) and growth rate (PD day^-1^) of U2OS cells untreated or treated with BIBR or 75 nM of TMOs, monitored weekly over 4 weeks. **D.** TRF analysis of genomic DNA extracted from U2OS cells untreated or treated with BIBR or 75 nM of TMOs.

