## Supplementary Material for "Thiomorpholino antisense oligonucleotides inhibit telomerase and limit cancer cell proliferation"

A

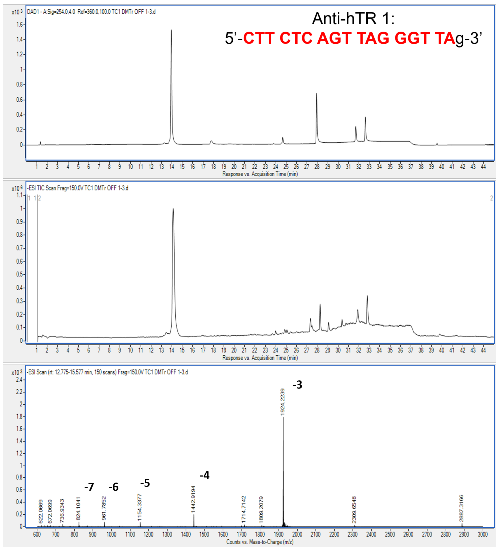

B

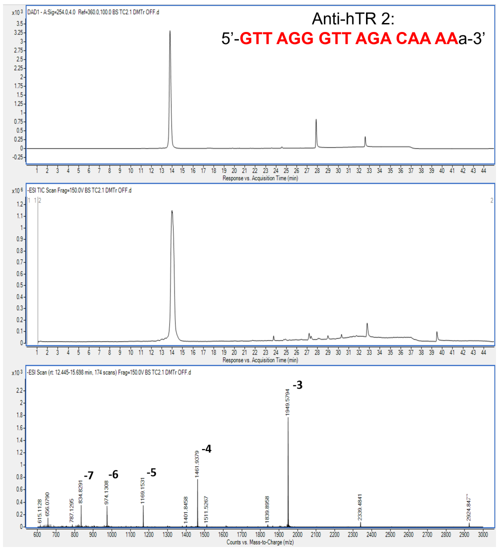

C

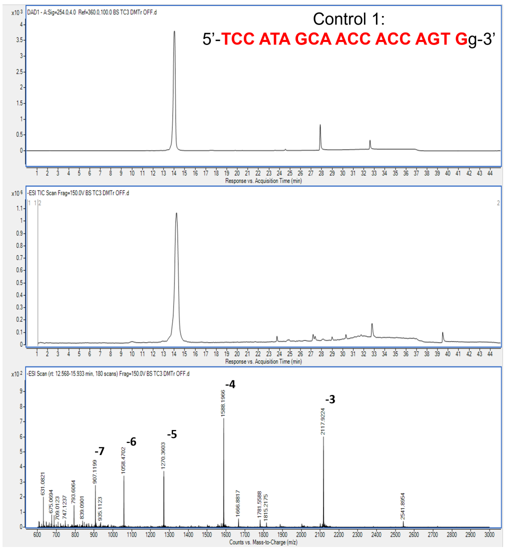

D

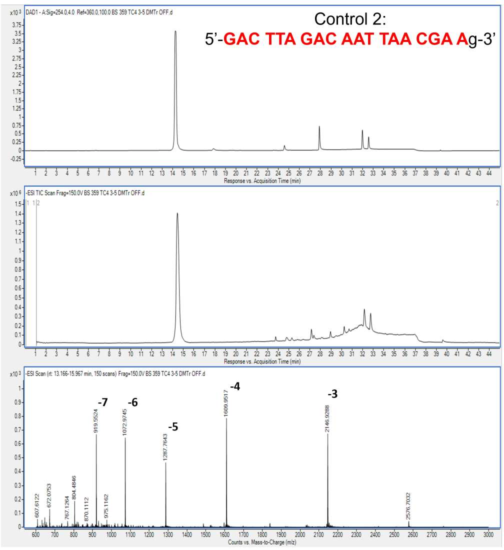

E

| TMO | Calculated MW | Measured MW |
|---|---|---|
| Anti-hTR1 | 5775.949 | 5775.672 |
| Anti-hTR2 | 5851.039 | 5851.738 |
| C1 | 6356.479 | 6356.767 |
| C2 | 6443.565 | 6443.786 |

F

| Duplex with RNA template+extensions | ΔTm (°C) |
|---|---|
| TMO-DNA anti-hTR1 | 10.0 ± 0.2 |
| TMO-DNA anti-hTR2 | 6.7 ± 0.3 |

G

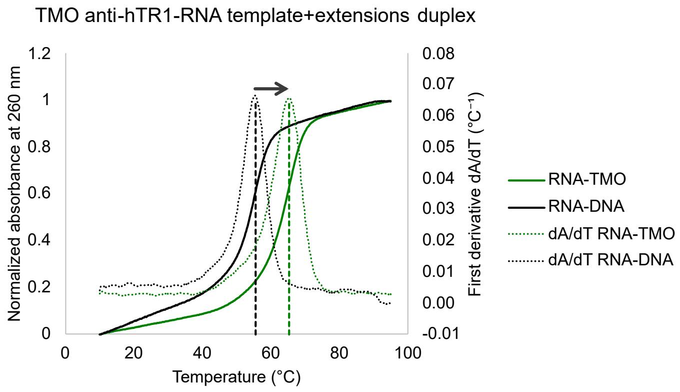

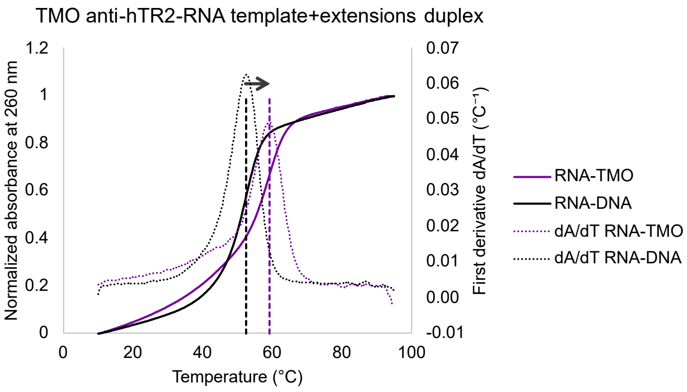

Supp. Figure 1

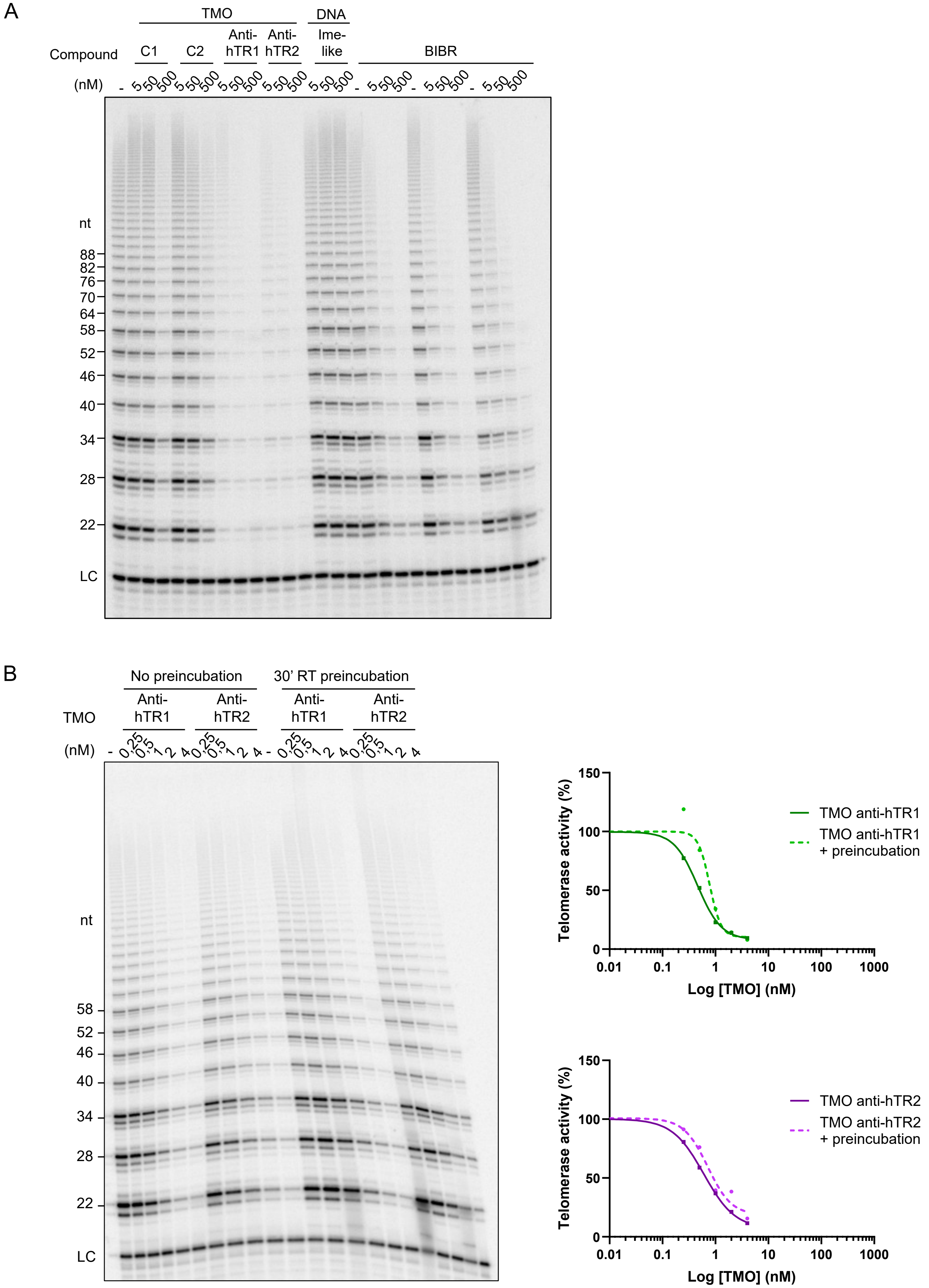

Supp. Figure 2

A

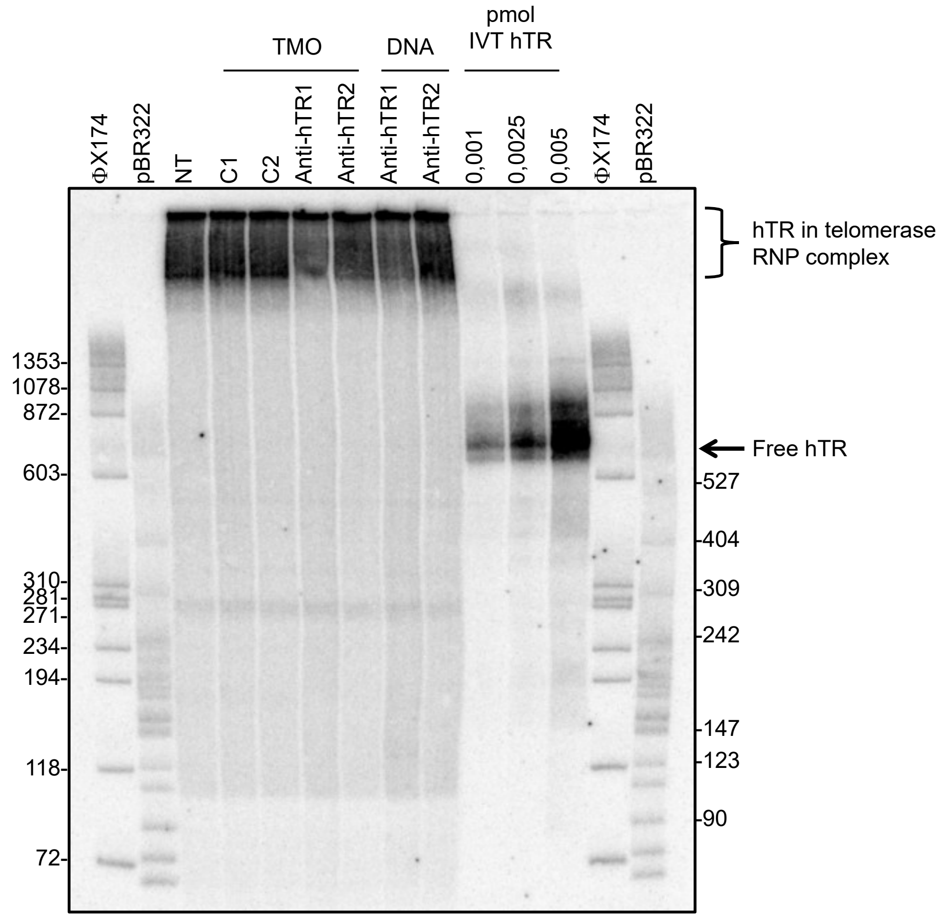

Supp. Figure 3

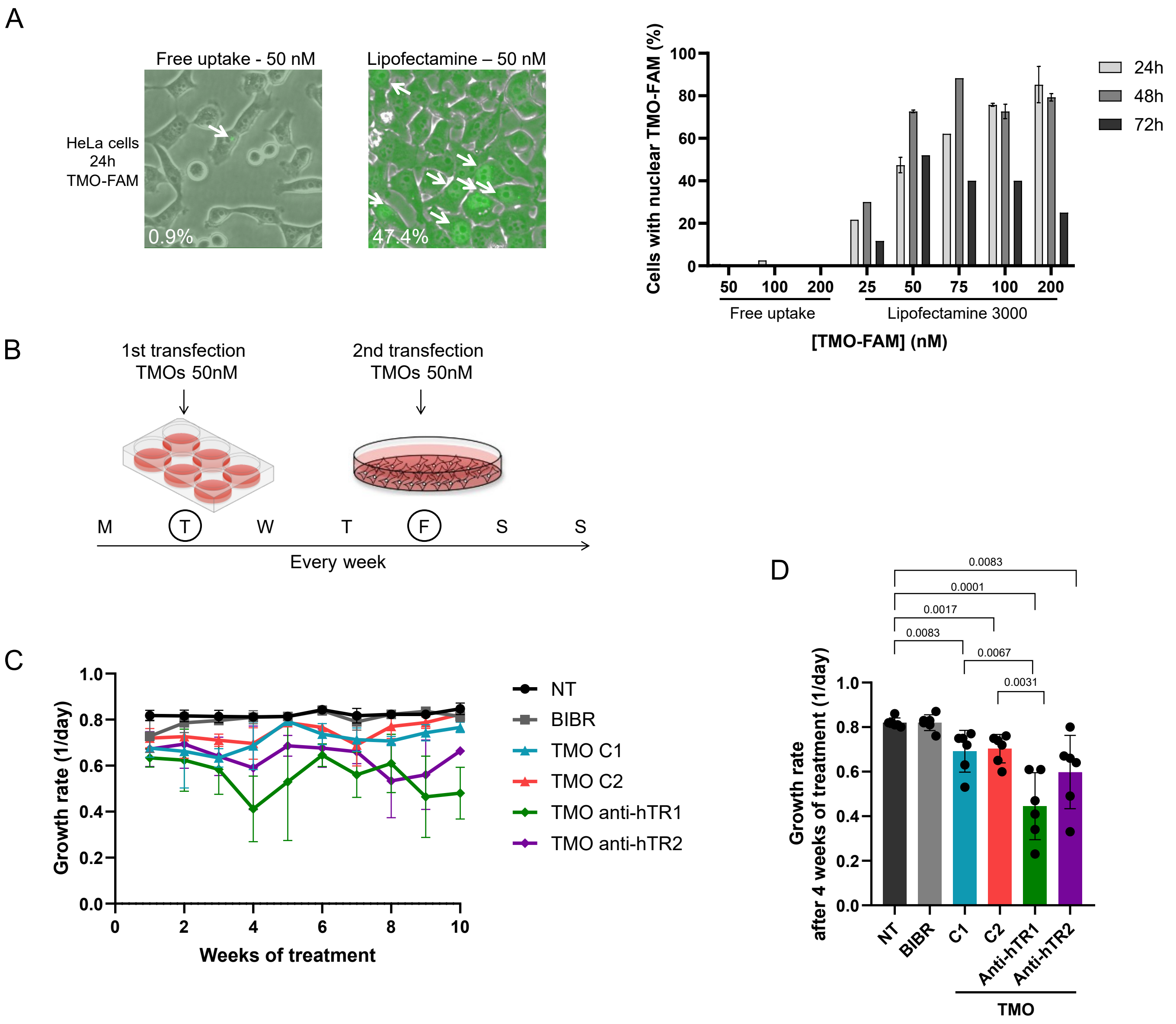

Supp. Figure 4

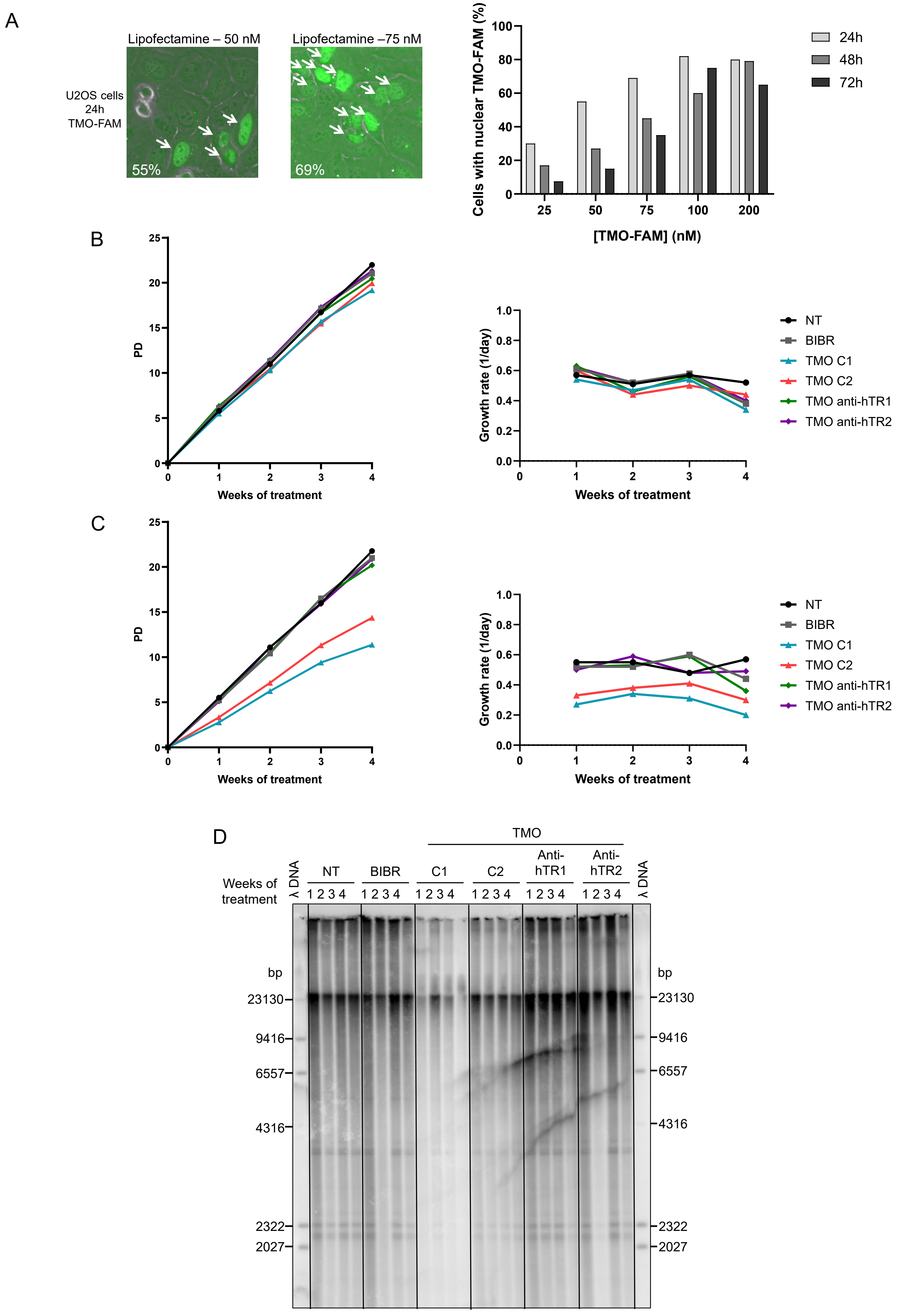

Supp. Figure 5

Supplemental_Table_S1: Oligonucleotides used in this study

| | |
|---|---|
| **TMOs** | |
| Anti-hTR1 | 5'-CTTCTCAGTTAGGGTTAG-3' |
| Anti-hTR2 | 5'-GTTAGGGTTAGACAAAAA-3' |
| Control 1 | 5'-TCCATAGCAACCACCAGTGG-3' |
| Control 2 | 5'-GACTTAGACAATTAACGAAG-3' |
| TMO-FAM | 5'-FAM-CCATAGCAACCACCAGTGGGGAG-3' |
| **DNA oligos** | |
| Anti-hTR1 | 5'-CTTCTCAGTTAGGGTTAG-3' |
| Anti-hTR2 | 5'-GTTAGGGTTAGACAAAAA-3' |
| Ime-like | 5'-TAGGGTTAGACAA-3' |
| hTR 130R-27 northern blot probe | 5'-CTTTTCCGCCCGCTGAAAGTCAGCGAG-3' |
| TelC TRF probe | 5'-TTAGGGTTAGGGTTAGGGTTAGGG-3' |
| **RNA oligos** | |
| 11-mer hTR template | 5'-CUAACCCUAAC-3' |
| 24-mer hTR template+flanking sequences | 5'-UUUUUUGUCUAACCCUAACUGAGA-3' |
